# Characterization and Preclinical Treatment of Rotational Force-Induced Brain Injury

**DOI:** 10.1101/2022.07.20.500670

**Authors:** Alan Umfress, Ayanabha Chakraborti, Suma Priya Sudarsana Devi, Raegan Adams, Daniel Epstein, Adriana Massicano, Anna Sorace, Sarbjit Singh, M. Iqbal Hossian, Shaida A. Andrabi, David K. Crossman, Nilesh Kumar, M. Shahid Mukhtar, Claire Simpson, Kathryn Abell, Matthew Stokes, Thorsten Wiederhold, Charles Rosen, Huiyang Luo, Hongbing Lu, Amarnath Natarajan, James A. Bibb

**Author notes:** Correspondence: James A. Bibb, University of Alabama at Birmingham, Department of Surgery, Center 1900 University Blvd, THT 1052 Birmingham, AL 35233.

## Abstract

Millions of traumatic brain injuries (TBIs) occur annually. TBIs commonly result from falls, traffic accidents, and sports-related injuries, all of which involve rotational acceleration/deceleration of the brain. During these injuries, the brain endures a multitude of primary insults including compression of brain tissue, damaged vasculature, and diffuse axonal injury. All of these deleterious effects can contribute to secondary brain ischemia, cellular death, and neuroinflammation that progress for weeks to months after injury and impede neurological recovery. While the linear effects of head trauma have been extensively modeled, less is known about how rotational injuries mediate neuronal damage following injury. Here, we developed a new model of rotational head trauma in rodents and extensively characterized the pathological, behavioral, and electrophysiological effects of rotational TBI (rTBI). We identify aberrant cyclin dependent kinase 5 (Cdk5) activity as a principal mediator of rTBI and show pharmacological inhibition of Cdk5 reduces the cognitive and pathological consequences of injury. Finally, we utilize Cdk5-enriched phosphoproteomics to uncover potential downstream mediators of rTBI. These studies contribute meaningfully to our understanding of the mechanisms of rTBI and how they may be effectively treated.

## INTRODUCTION

Traumatic brain injuries occur in epidemic proportions with over 1.5 million injuries occurring annually in the United States alone (Capizzi et al., 2020). During a TBI, the brain undergoes both linear and rotational forces, compressing tissue and damaging axonal connections. While the linear effects of injury primarily result in focal brain damage, rotational acceleration of the brain produces shearing forces resulting in both focal and diffuse neuropathology. Diffuse axonal shearing as the result of rotational TBI (rTBI) causes persistent neuropathological consequences of brain trauma (Guskiewicz et al., 2007, Mihalik et al., 2007, Zhang et al., 2001). At the time of injury, rapid brain acceleration/deceleration creates a disruption of brain tissue resulting in contusion, blood vessel damage, hemorrhage, and axonal shearing (McKee and Daneshvar, 2015). TBI results in acute dysfunctions in cognition including loss of consciousness, headaches, and vison problems (Riggio and Wong, 2009). Secondary damage evolves over the following days, months, and lifetime of survivors leading to impairments in memory and motor function, anxiodepressive disorders, and neurodegeneration (Pavlovic et al., 2019, Arciniegas et al., 2002, Cuthbert et al., 2015, Adams et al., 2018).

At cellular resolution, rTBI induces axonal shearing resulting in massive neuronal depolarization and ionic influx (Nortje and Menon, 2004, Masel and DeWitt, 2010, Davis, 2000, Povlishock and Christman, 1995, Raghupathi, 2004). In response, activation of voltage-gated Ca^2+^ channels induce excitotoxic release of glutamate. Following excitotoxicity, cerebral edema, oxidative stress, and cellular death all contribute to the acute phase of injury. (Capizzi et al., 2020, Chen et al., 2019, Greve and Zink, 2009). After initial trauma, a delayed spreading process of injury occurs. The injured brain exhibits increased sensitivity to secondary ischemic insult and persistent excitotoxicity (Shi et al., 2019).

The disruption of intracellular signaling cascades serves as a convergence point of ischemia, inflammation, and excitotoxicity where the activation of downstream effectors induces cellular demise. One such excitotoxic pathway is the dysregulation of the protein kinase cyclin dependent kinase 5 (Cdk5) via the calpain family of Ca^2+^-activated neutral proteases (Yousuf et al., 2016). Following cellular damage, activated calpain protease cleaves the coactivator of Cdk5, p35, into a truncated aberrant coactivator, p25 (Kusakawa et al., 2000). Conversion of Cdk5/p35 to Cdk5/p25 by calpain confers neurotoxic activity upon the kinase, resulting in neuronal injury and death (Meyer et al., 2008). Aberrant Cdk5/p25 contributes to virtually all neurodegenerative diseases and is key to the general processes by which neurotoxicity occurs (Barnett and Bibb, 2011, Torres-Altoro et al., 2011, Meyer et al., 2008). Conditional knockout of Cdk5 confers neuroprotection from cortical impact, ischemic stroke, and mouse models of Alzheimer’s Disease (AD) (Meyer et al., 2014b, Yousuf et al., 2016, Camins et al., 2006). Therefore, we hypothesized that aberrant Cdk5/p25 activity may mediate the neuropathological and neurocognitive effects of repeated and rotational brain injuries. To test this, we designed and characterized a novel model of rTBI and assessed the neuropathological effects of injury in the acute and subacute phases, including aberrant Cdk5/p25 activation. We assessed a new systemic Cdk5 inhibitor (25-106) as a potential treatment for rTBI. Finally, we conducted Cdk5-enriched phosphoproteomics to identify novel downstream effectors of brain injury. Together, these studies provide a better understanding of the mechanisms mediating injury by rTBI and point to a potentially effective therapeutic approach.

## RESULTS

### Development and characterization of a rotational Traumatic Brain Injury (rTBI) model in rats

To assess the negative consequences of rotational head injury, we developed a novel rTBI model with the ability to impart clinically relevant angular accelerations (Figure 1A). The model consists of a pneumatic-chain driven pendulum arm attached to an animal carrier. This pendulum rotates about a horizontal shaft. The rodent is placed in a restraint inserted onto a fixed mount at the end of the pendulum (Figure 1B, Figure 1 Supplement 1A) and attached to the freely rotating helmet assembly bolted to the restraint mount (Figure 1C), allowing pure coronal plane head rotation. A ventral strike plate (Figure 1D, Figure 1 Supplement 1A) transfers impact momentum between the harness and impact fixture, imparting rotational force on the subject’s head. The pendulum is rapidly propelled forward using a compressed N_2_-driven 2-way solenoid valve (Figure 1E,F) that subsequently drives an air motor inducing rapid chain rotation. Accelerometry is derived from a helmet mounted inertial measurement unit (IMU) model 633, a 6-degrees of freedom transducer incorporating accelerometers and gyroscopes comprised of micro-machined silicon sensors (Figure 1 Supplement 1B). IMU data is captured in LabVIEW and converted to acceleration measured in gravitational constants (g values), which are converted to angular acceleration (rad/sec^2^).

**Figure 1.**
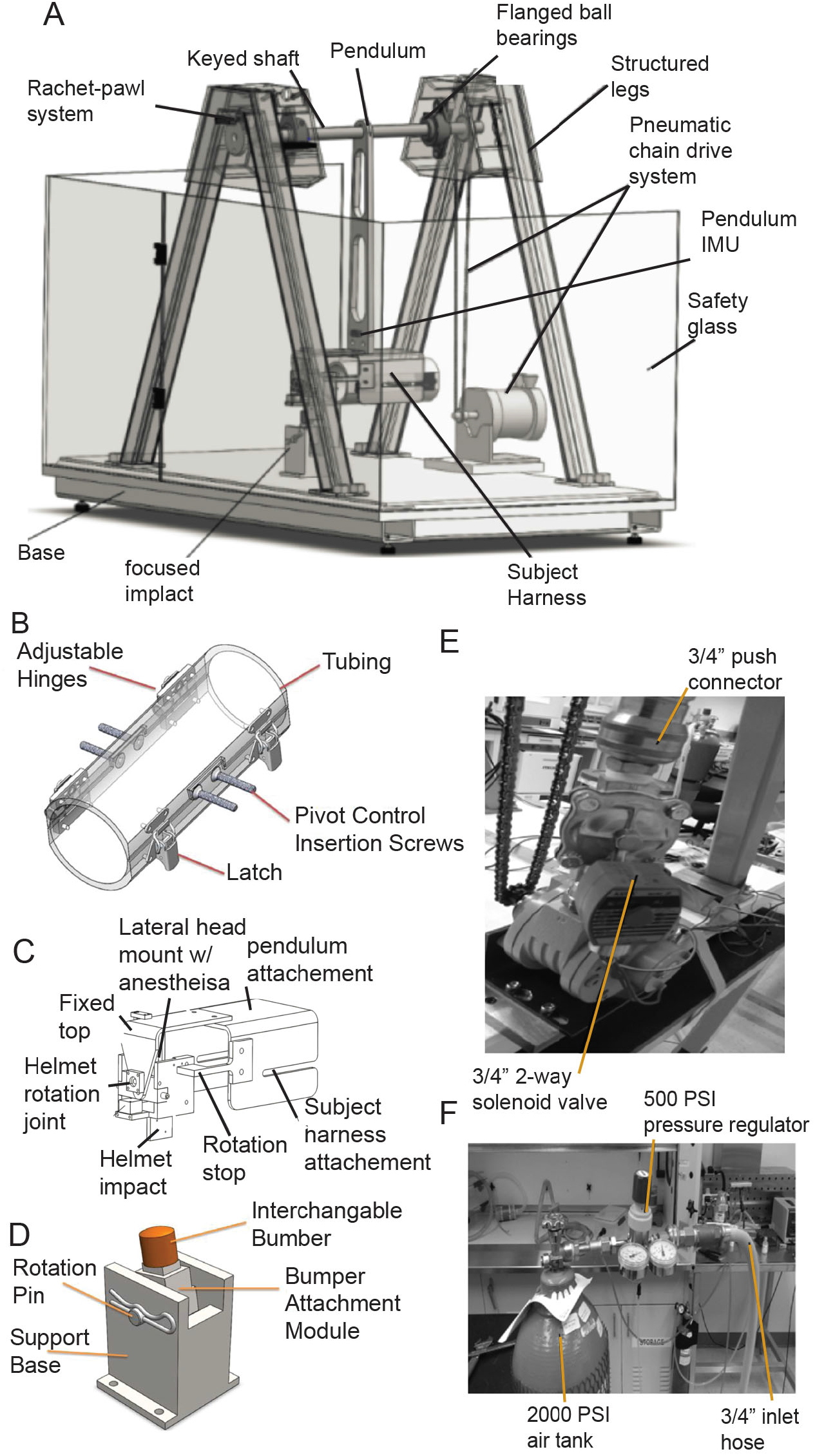
Novel rodent model of rotational traumatic brain injury (rTBI) A. Rotational head injury model, components, and complete design. B. Animal restraint. C. Helmet and subject constraint mount. D. Ventral Strike plate. E. Pneumatic-chain drive system. F. Pneumatic pressure system.

To validate helmet sensor accelerometry, we utilized high speed video telemetry to determine angular speed of the helmet and compared these values to those derived from the helmet sensor (Figure Supplement 1C-F). Video telemetry and helmet sensor showed highly overlapping determinates for angular speed. Typical machine runs induce average helmet peak rotational acceleration outputs in LabView averaging 482Krad/s^2^ with limited variability across 5 sampling runs of 14.15 Krad/s^2^ (Figure Supplement 1G). These angular accelerations are consistent with human to animal scaling laws for mild brain injuries (Rowson et al., 2009, Viano et al., 2009), suggesting rTBI studies using this model in rodents may readily translate to human conditions.

### Neuropathological consequences of rTBI

The pathological consequences of rTBI include damaged vasculature, diffuse axonal injury, inflammation, and cellular demise (McKee and Daneshvar, 2015). To investigate the effects of rTBI, we first conducted PET/CT neuropathology studies. For these studies, we made use of a diagnostic molecular probe in current clinical use for the detection of the neuroinflammatory marker, translocator protein (TSPO). This probe, [^18^F]DPA-714 detects activated microglia resulting from neuroinflammation in neurogenerative and neuroinflammatory conditions (Coughlin et al., 2015, Israel et al., 2016, Werry et al., 2019). TSPO imaging revealed differential uptake diffusely throughout the brains of both control and rTBI rats 7 days post-injury (Figure 2A). Interestingly, standard uptake values (SUV) were significantly increased in several brain regions including amygdala, hippocampus CA1 layer, and the basolateral amygdaloid nucleus of rTBI rats 7 days post-injury in comparison to controls (Figure 2B-D). Effects were also observed diffusely throughout whole cortex with cortical areas including perirhinal and primary somatosensory cortex showing higher labeling following rTBI (Figure 2 Supplement 1A-C). The differential labeling was region-specific as no effect was detectible when signal throughout whole brain was quantitated (Figure 2 Supplement 1D). Also, regions such as hippocampus layer CA3 and cerebellum exhibited no change in response to impact (Figure 2 Supplement 1E, F). Thus, rTBI caused brain region specific alterations in this biomarker of neuroinflammation, consistent with effects observed in humans (Coughlin et al., 2015).

**Figure 2.**
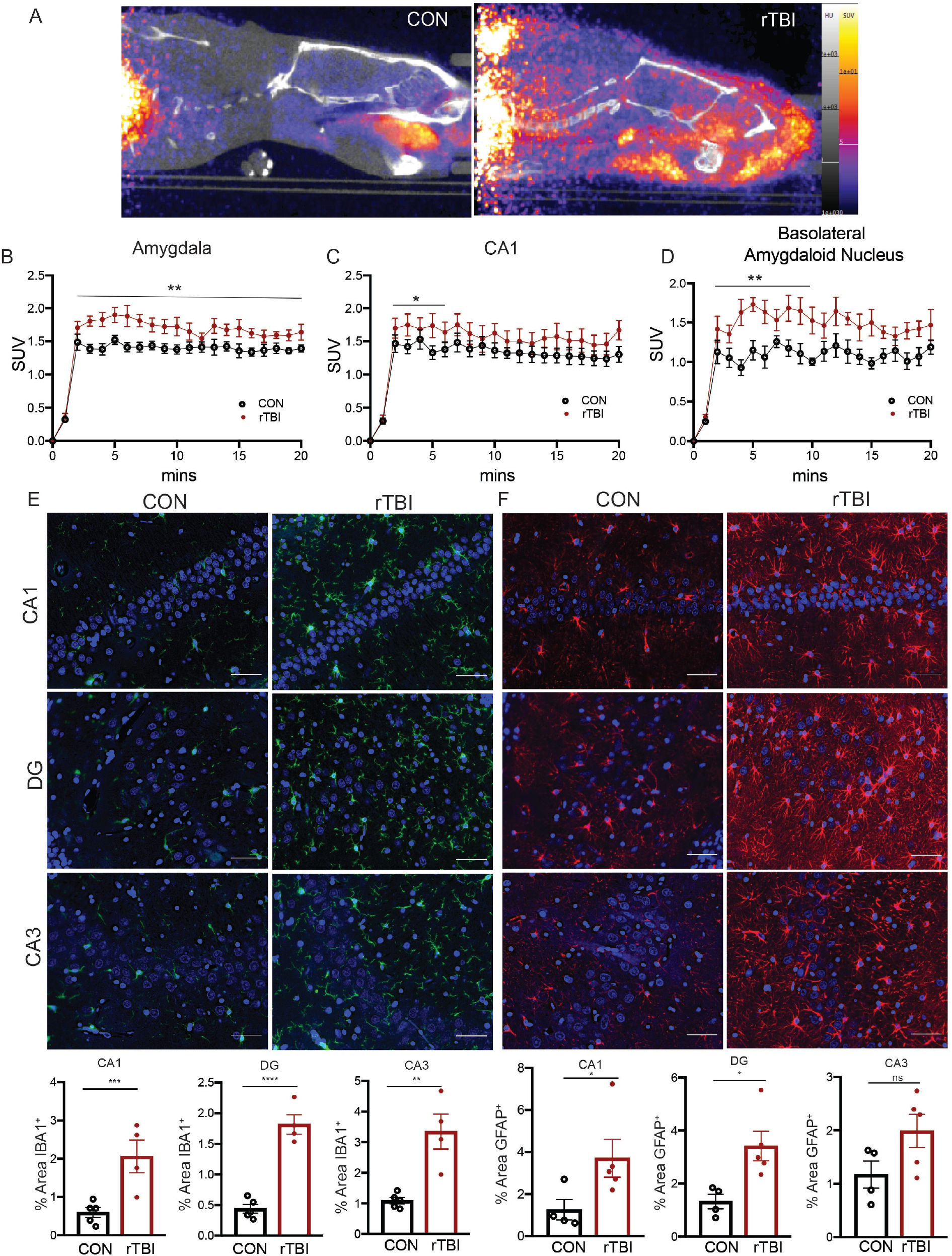
Neuropathological effects of rTBI. A. Representative PET/CT images for TSPO uptake in control (left) and rTBI (right) rats. B. Quantitative Standard uptake values (SUV) for TSPO radioligand in amygdala (Time: F(20,200) = 110.6, p < 0.0001; Treatment: (1, 10) = 7.771, p = 0.0192; Interaction: (20, 200) = 2.321, p = 0.0017) two-way-RM ANOVA C. CA1 (Time: F(5,50) = 216.4, p < 0.0001; Treatment: F (1,10) = 1.711, p = 0.2202; Interaction: F(5,50) = 2.614, p = 0.0355) two-way-RM ANOVA D. Basolateral amygdaloid nucleus (Time: F(10,100) = 4 9.67, p < 0.0001; Treatment: F(1,10) = 13.97, p=0.0039; Interaction: F(10,100) = 2.605, p = 0.0075). E. Histological staining and quantitation for IBA1^+^ microglia within CA1, p = 0.0083 Student’s *t*-test; DG, p = Student’s *t*-test; and CA3, p = 0.0032 Student’s *t*-test. F. Histological staining and quantitation for GFAP^+^ astrocytes within CA1, p = 0.0317 Mann Whitney test (p = 0.0300, Shapiro-Wilk); DG, p = 0.0174 Student’s *t*-test; and CA3, p = 0.0878 Student’s *t*-test. All data are means ± SEM, *p < 0.05, **< 0.01, **< 0.01, ****< 0.001. Scale bars = 50 μm.

In addition to *in vivo* imaging, immunostaining was used to assess neuroinflammatory effects of rTBI. For these experiments, layers of the hippocampal formation were examined, as rTBI effects within this brain region in humans have been linked to consequent memory impairments (Arulsamy et al., 2018, Atkins, 2011, Bigler et al., 2002). Indeed, broad increases in microgliosis were detected throughout the CA1, dentate gyrus (DG), and CA3 subfields of the hippocampus (Figure 2E) 48 h post-injury. Furthermore, rats subjected to rTBI showed increased astrogliosis throughout the CA1 and dentate gyrus (DG), with a trend also toward increases in CA3 region of the hippocampus (Figure 2F). Together, these *in vivo* imaging and histological studies demonstrate rotational head injury induces neuroinflammation in acute and sub-acute phases of injury.

### Behavioral and Neurophysiological consequences of rTBI

Persistent memory impairment is a common outcome following TBI (Arciniegas et al., 2002). Patients report symptoms including both anterograde and retrograde amnesia, as well as, cognitive deficits in attention, processing speed, and executive functioning (Rabinowitz and Levin, 2014). These deficits often correlate with structural damage to temporal brain areas (Arulsamy et al., 2018) and damage within the hippocampal formation following TBI can contribute to long-lasting deficits in memory (Mathias and Mansfield, 2005). Therefore, we investigated the functional and physiological consequences of rTBI on hippocampal-dependent memory and plasticity in our model. To assess effects on contextual and cued fear learning and memory, rats (7 days post-injury) were placed in a novel context, analyzed for baseline freezing responses, and subsequently exposed to fear conditioning cue/foot shock pairings for assessment of learning and memory function (Figure 3A). Rats subjected to rTBI displayed no alteration in baseline freezing rates. However, rTBI rats showed a marked decrease in freezing behavior in response to re-exposure to the adverse context. Additionally, rTBI rats displayed a reduction in cue-induced freezing rates in a novel context, suggesting that both hippocampal- and amygdala-mediated components of this learned behavior were impaired. This reduction in freezing responses was not confounded by injury-induced damage to nociceptive circuitry, as shock stimulus pain thresholds for animal flinching, jumping, and vocalizing pain were not altered by rTBI (Figure 3B). Thus, rTBI caused learning and memory deficits that persisting into the sub-acute phase of injury.

**Figure 3.**
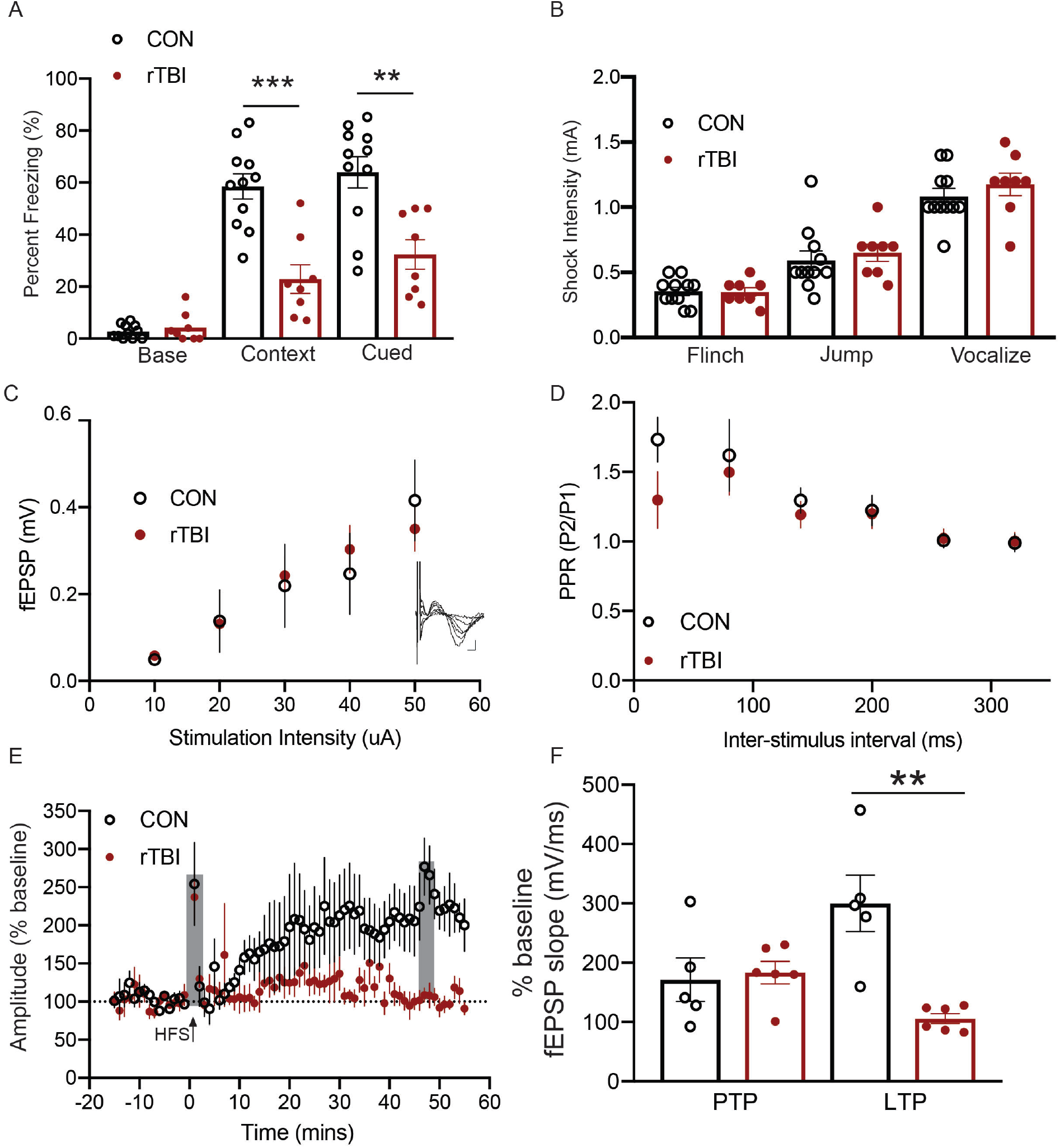
Behavioral and neurophysiological consequences of rTBI. A. Fear conditioning freezing rates for baseline, Contextual, p = 0.0001 Student’s *t*-test, and Cued, p = 0.0018 Student’s *t*-test fear learning. B. Shock sensitivity thresholds to flinch, jump, or vocalize pain. C. Input-Output curve of CA3-CA1 fEPSP recordings (Inset: individual traces for each of the stimulus intensities). D. Paired Pulse Ratio (PPR) across inter-stimulus interval. E. Assessment of the effect of rTBI on hippocampal plasticity after HFS. (PTP outlined grey 0-2 min, LTP outlined grey 45-55 min). F. Summary of PTP and LTP fEPSP slopes, p= 0.7662, PTP; p = 0.0017, LTP Student’s *t*-test. All data are means ± SEM, *p < 0.05, **< 0.01, ***< 0.001.

The observed mnemonic impairments in response to rTBI suggest that hippocampal circuitry function may be damaged. To investigate this possibility, field excitatory post synaptic potentials (fEPSP) recordings were taken from hippocampal CA3-CA1 circuitry (7 days postinjury) (Figure 3C-F). Synaptic excitability, as assessed via stimulus-to-fEPSP amplitude ratios (I/V curves), was unaffected by rTBI (Figure 3C). The paired-pulse ratio paradigm was used to assess changes on neurotransmitter release. Similarly, this metric of short-term plasticity remained unaltered in rTBI rats (Figure 3D). Additionally, high frequency stimulation (HFS) induced a 171.4 ± 36% post-tetanic potentiation (PTP) in control rats compared to baseline (Figure 3E, F). This effect was similar (183.4 ± 18%) in rTBI brains. HFS also induced robust (300 ± 47%) long-term potentiation (LTP) of fEPSP slope compared to baseline 45 min poststimulus at Schaffer collateral-CA1 synapses of control mice. (Figure 3E, F). In contrast, tetanic stimulation caused a markedly smaller effect (106 ± 8%) in rTBI rats, almost completely ablating the ability to induce LTP. Thus, rTBI caused a notable impairment in hippocampal long-term synaptic plasticity as a likely mechanism underlying the deficit it induced in learning and memory.

### Excitotoxicity and rTBI evoke aberrant Cdk5 activity and Cdk5 inhibition is neuroprotective

TBI shares mechanistic features with other forms of neuronal injury in that excitotoxicity triggers loss of Ca^2+^ homeostasis (Blennow et al., 2012). The activation of calpain and consequent truncation of the Cdk5 coactivator p35 to its aberrant coactivator p25 has been implicated in various forms of neuronal injury including ischemic stroke, cortical impact, and blast TBI (Meyer et al., 2014b, Yousuf et al., 2016, Hernandez et al., 2018). Generation of the aberrantly active Cdk5/p25 complex is neurotoxic (Patrick et al., 1999). Cdk5/p25 hyperphosphorylates tau in models of AD and causes neuronal cell death (Cruz et al., 2003). Thus, aberrant Cdk5 invoked by excitotoxicity is a common neuronal injury mechanism shared by many neuropathological conditions (Meyer et al., 2014b, Pao and Tsai, 2021).

To better understand the role of aberrant Cdk5 in mediating excitotoxicity in rTBI, we first treated *ex vivo* brain slices with high concentrations of NMDA (100 μM) and glycine (gly) (50 μM) or subjected acutely prepared brain slices to oxygen and glucose deprivation (OGD) to induce ischemic excitotoxicity (Figure 4 Supplement 1). Each of these treatments caused significant generation of p25 (Figure 4 Supplement 1A,B). In comparison with *ex vivo* conditions, rTBI induced comparable increases in p25 production *in vivo* (Figure 4A). This effect corresponds to the excitotoxic activation of calpain as demonstrated by the breakdown of the calpain reporter, Fodrin into cleaved 150 and 120 kDa products 48 h post-injury (Figure 4B) (Czogalla and Sikorski, 2005). Thus rTBI, like other forms of excitotoxic neuronal injury activates the Ca^2+^-calpain-Cdk5/p25 neurotoxicity pathway as a potential mechanism mediating loss of neuronal function.

**Figure 4.**
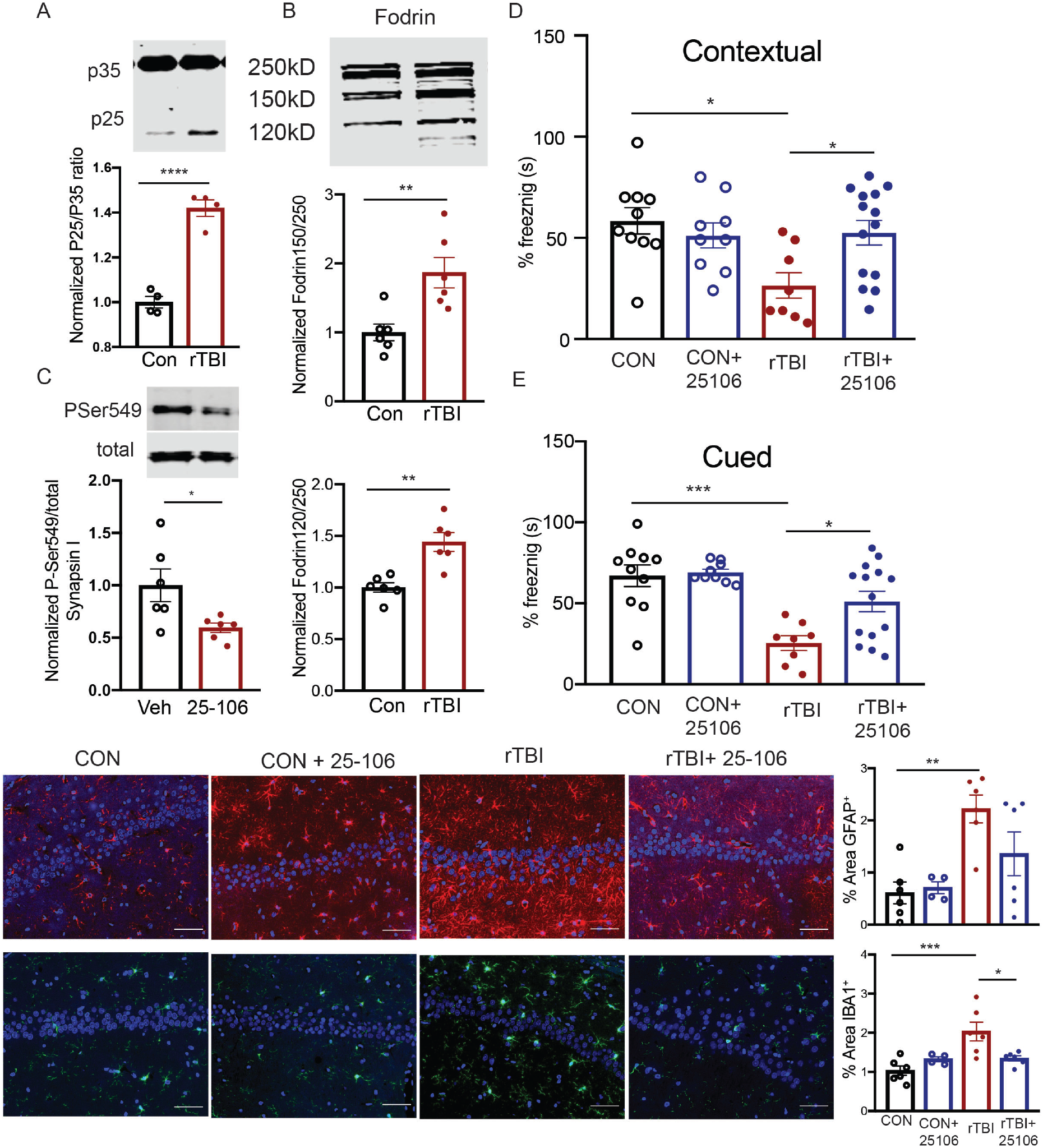
rTBI induces aberrant activation of Cdk5 and inhibition is neuroprotective. A. Quantitative immunoblot of P25 generation following rTBI, p <0.0001 Student’s *t*-test. B. Quantitative immunoblot of Fodrin breakdown products via calpain, p = 0.0062 (top), p = 0.0014 (bottom) Student’s *t*-test. C. *In vivo* inhibition of Cdk5 assessed via quantitative immunoblot for P-Ser549/total Synapsin I after treatment with 50 mg/kg 25-106, p = 0.0305 Student’s *t*-test. D. Fear conditioning freezing rates in contextual conditioning after rTBI treatment with 25-106 F(3,37) = 4.057, p = 0.0137, ANOVA; (Con-TBI p = 0.0132, TBI-TBI + 25106, p =0. 0384 Holm-Sidak *post hoc*). E. Fear conditioning freezing rates in cued conditioning after rTBI treatment with 25-106 F(3,37)= 10.07, p < 0.0001 ANOVA; (Con-TBI p = 0.0002, TBI-TBI + 25106 p = 0.0137 Holm-Sidak *post hoc*). F. Immunohistochemistry and quantitation of GFAP^+^ astrocytes in CA1 following rTBI and treatment with 25-106 F(3,18) = 6.295, p = 0.0041 ANOVA; (Con-TBI, p = 0.0054, Con-TBI + 25106, p = 0.2259, Holm-Sidak *post hoc*). G. Immunohistochemistry and quantitation of IBA1^+^ microglia in CA1 following rTBI and treatment with 25-106 F(3,18) = 8.070, p = 0.0013 ANOVA; (Con-TBI, p = 0.0009, TBI-TBI+ 25106 p = 0.0209, Holm-Sidak *post hoc*). All data are means ± SEM, *p < 0.05, **< 0.01, ***< 0.01. Scale bars = 50 μm.

While over-expression of p25 is neurotoxic (Meyer et al., 2008, Cruz et al., 2003), pharmacological inhibition of Cdk5/p25 activity is neuroprotective *in vitro* (Meyer et al., 2014b). Furthermore, conditional knockout of Cdk5 is neuroprotective *in vivo* (Meyer et al., 2014b, Yousuf et al., 2016). However, effective targeting of Cdk5 as a post-insult therapeutic approach has not been possible due to the lack of brain penetrant Cdk5/p25 inhibitors. Recently, we discovered a brain diffusible Cdk5 inhibitor (25-106) with exponential brain distribution kinetics and 30-fold greater specificity for Cdk5 over other cyclin dependent kinase family members (Umfress et al., 2022). We hypothesized that this inhibitor could have rTBI therapeutic potential by blocking aberrant Cdk5/p25 activity. We first verified that this compound acted as a Cdk5 inhibitor in brain tissue by assessing the effect of slice treatment with 25-106 on phosphorylation of the Cdk5 reporter, Thr75 DARPP32 (Figure 4 Supplement 1C). Indeed, this site was strongly attenuated in brain slices by 25-106 treatment. Next, we assessed the neuroprotective capacity of 25-106 under excitotoxic conditions. First, primary neuronal cultures were subjected to OGD which reduced neuronal viability (Figure 4 Supplement 1D). This effect was attenuated in cells first treated with 25-106. Similarly, in *ex vivo* brain slices, OGD induced cellular death. This response was significantly reduced by incubating brain slices with 25-106 (Figure 4 Supplement 1E). Additionally, the neuroprotective effect following Cdk5 inhibition was observed in *ex vivo* brain slices treated with high concentrations of NMDA/Gly (Figure 4 Supplement 1F). These data confirm 25-106 as an inhibitor of Cdk5 in brain tissue and suggest it is neuroprotective in *in vitro* and *ex vivo* models of excitotoxicity.

To test the *in vivo* efficacy of 25-106, rats were subjected to rTBI and treated with 25-106 within 10 min after undergoing the injury procedure. Cognitive recovery was then assessed by fear conditioning 7 days post-injury (Figure 4D,E). As previously observed, rats subjected to rTBI showed significant reductions in contextual fear learning and memory (freezing behavior). Also, treatment with 25-106 in the absence of rTBI had no effect on basal memory performance. In contrast, rTBI rats treated with 25-106 displayed no significant memory impairment in comparison to uninjured control animals and were significantly improved in mnemonic function compared to animals that received rTBI without Cdk5 inhibition (Figure 4D). Similarly, rats subjected to rTBI alone showed significant reductions in cued fear learning and memory, while rTBI rats treated with 25-106 displayed cued-leaning and memory rates similar to control animals and significantly improved from rTBI alone (Figure 4E). Thus, acute post-injury Cdk5 inhibition almost completely blocked cognitive impairment in response to rTBI.

To assess the ability of 25-106 to ameliorate the neuropathological outcomes associated with impaired hippocampal memory, neuroinflammation was assessed via staining for astrogliosis and microgliosis 14 days post-injury (Figure 4F,G). While rTBI animals alone showed the significant increases in CA1 astrocyte reactivity (GFAP^+^), similar to that observed at 48 h post-rTBI, 25-106 treatment attenuated this effect. Similarly, 25-106 prevented the neuroinflammatory activation of microglia in the hippocampus CA1 layers (Figure 4G). In addition to these neuroprotective effects in the CA1 region, increases in gliosis occurred in other hippocampal layers including the dentate gyrus, and CA3 following rTBI and was attenuated by 25-106 (Figure 4 Supplement 2 A-F). Altogether, these data demonstrate the therapeutic potential of systemic Cdk5 inhibition via 25-106 to neuroprotect from rTBI.

### Phosphoproteomic alterations following rTBI

To investigate the downstream effectors of Cdk5/p25 that may mediate neuronal demise and subsequent cognitive impairments associated with injury, we conducted Cdk5 phosphorylation site-directed phosphoproteomics and global proteomics of the hippocampus from control, rTBI, and rTBI rats treated with 25-106 (rTBI + 25-106) 48h post-injury. Robust changes in the Cdk5 phospho-landscape following rTBI and rTBI+ treatment with 25-106 were apparent with 3,021 uniquely modified sites detected via LC/MS-MS (Figure 5A-C). Rotational head trauma resulted in 653 upregulated phospho-sites with a fold change (FC) ≥1.3 (Figure 5A), and a corresponding decrease in 650 ≤-1.3 FC phospho-sites. Interestingly, differentially modified sites included alterations in the phosphorylation of proteins associated with AD such as Microtubule Associated Protein 2 (MAP2), Spinocerebellar Ataxia (SCA) associated protein, and Diacylglycerol Lipase Alpha (DAGLA) (Iqbal et al., 2005, Carlson et al., 2009). Additionally, the potential Cdk5-dependent phosphorylation states of proteins associated with cellular demise such as, apoptosis-stimulating protein of p53 (PPP1R13B), and homeostatic synaptic excitability such as GRIN2a (GluN2A) were altered by rTBI (Figure 5A) (Samuels-Lev et al., 2001, Yong et al., 2021).

**Figure 5.**
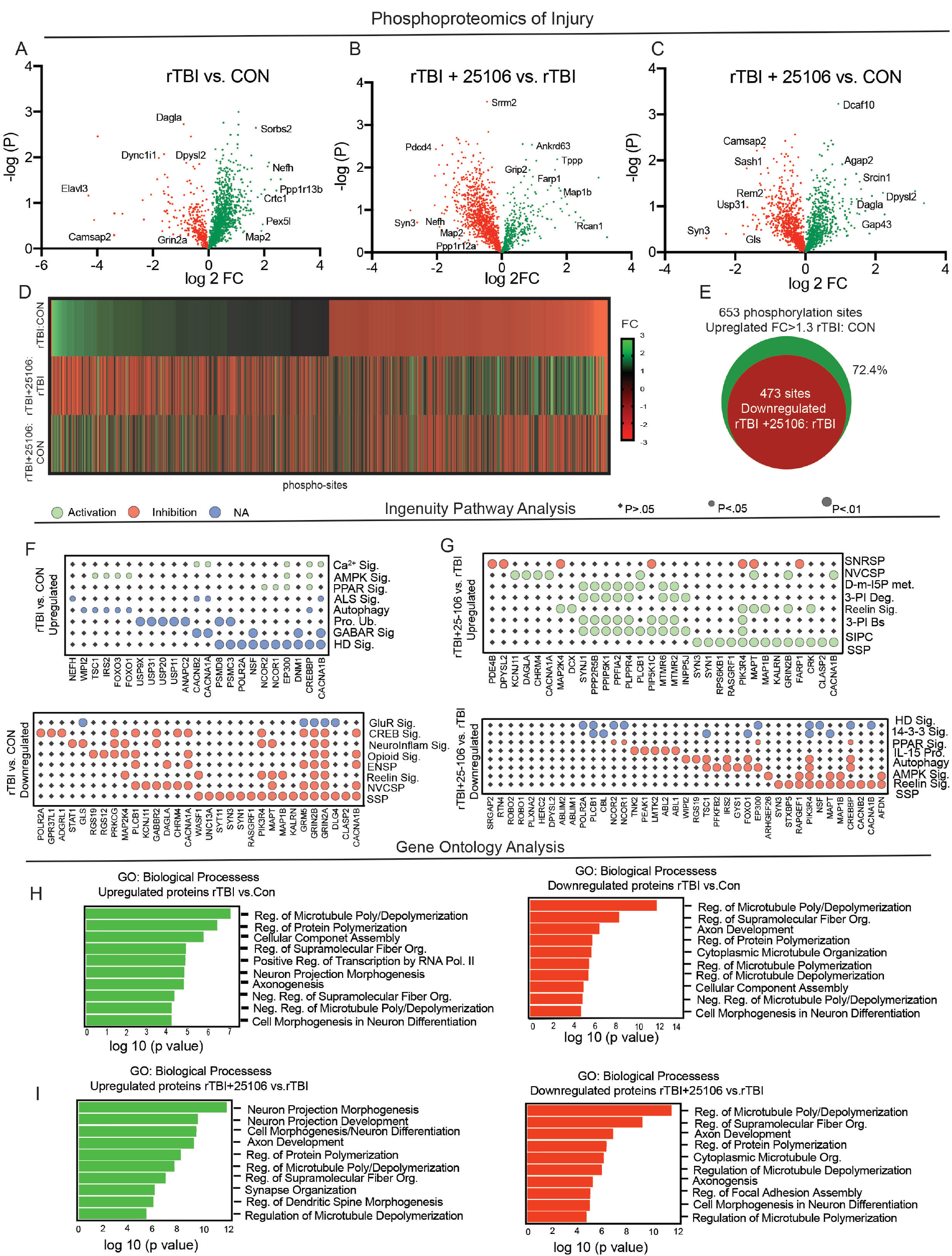
Cdk5-enriched phosphoproteomics of rTBI. A. Volcano plot of differentially phosphorylated proteins between rTBI and con rats (increased sites in green decreased sites in red). B. Volcano plot of differentially phosphorylated proteins between rTBI + 25-106 and rTBI alone rats. C. Volcano plot of differentially phosphorylated proteins between rTBI + 25-106 and con rats. D. Heatmap of all differentially phosphorylated sites across three comparisons in A-C expressed by fold change of each phosphor-site E. Venn diagram representing overlapping sites with positive FC ≥ 1.3 in rTBI:CON and negative FC in rTBI + 25-106:rTBI. F. Dot-plot display of Ingenuity Pathway Analysis of phospho-proteins and canonical pathways upregulated FC ≥ 1.3 by rTBI:Con (Top) and downregulated with a FC ≤ −1.3 by rTBI:Con (bottom). G. Dotplot display of Ingenuity Pathway Analysis of phospho-proteins and canonical pathways upregulated FC ≥ 1.3 by rTBI + 25106:rTBI (Top) and downregulated with a FC ≤ −1.3 by rTBI + 25-106:rTBI (bottom). H. Gene ontology of phosphoproteins biological processes upregulated FC ≥ 1.3 (left, green) or downregulated FC ≤ −1.3 (right, red) in rTBI:Con. I. Gene ontology of phosphoproteins biological processes upregulated FC≥1.3 (left, green) or downregulated FC≤-1.3 (right, red) in rTBI + 25-106:rTBI

We also observed a reciprocal relationship between the Cdk5 consensus site phospho-landscape of rTBI rats in comparison to rTBI rats treated with 25-106 (Figure 5D,E), as many phospho-sites that were increased by rTBI were reduced in rTBI rats treated with 25-106. Specifically, of the 653 phospho-sites upregulated by rTBI, 473 of these sites were decreased in the hippocampus of rTBI rats treated with 25-106 representing an inverse relationship of 72.4% of the phospho-sites (Figure 5E).

Ingenuity Pathway Analysis was used to identify key molecules and interacting pathways evoked by injury (Figure 5F,G). Upregulated pathways from rTBI highlight molecules linked to neuronal impairments. These included altered phosphorylation states of molecules involved in Ca^2+^ signaling (Ca^2+^ Sig.), AMPK signaling (AMPK Sig.), GABA receptor signaling (GABAR Sig.), and protein ubiquitination (Pro. Ub.), as well as pathways invoked in neurodegeneration including amyotrophic lateral sclerosis signaling (ALS Sig) and Huntington’s disease signaling (HD Sig.) (Figure 5F, top). Pathways assessed for proteins in which Cdk5-dependent phosphorylation states decreased following rTBI include glutamate receptor signaling (GluR Sig.), CREB signaling, reelin signaling, and synaptogenesis signaling pathways (SSP), all part of networks necessary for homeostatic neuronal function (Figure 5F, bottom). Furthermore, the reduced phospho-states following rTBI evoked alterations in molecules associated with neuroinflammatory signaling (Neuroinflam Sig.) and neurovascular coupling signaling pathways (NVCSP) (Figure 5F). Interestingly, rTBI rats treated with 25-106 (rTBI + 25106) invoked phosphorylation state increases in pathways such as neurovascular coupling signaling pathway (NVCSP) and synaptogenesis signaling pathways (SSP), both of which displayed a reciprocal relationship in untreated rTBI rats (Figure 5G). Rats subjected to rTBI+25-106 also showed many phosphorylation state-dependent increases in proteins associated with inositol phosphate signaling including D-myo-inositol-5-phosphate metabolism (D-m-IP5 met.), 3-phosphoinsitide biosynthesis (3-PI Bs) and degradation (3-PI Deg.), as well as, the super pathway of inositol phosphate compounds (SIPC) (Figure 5G).

Additionally, we observed an inverse relationship in the pathways implicated in the phospho-sites that decreased following rTBI + 25-106. Here, we observed decreased phosphorylation of proteins associated with Huntington’s disease signaling (HD Sig.), autophagy, and AMPK signaling, all of which were upregulated by rTBI alone (Figure 5G). Also, rTBI + 25-106 decreased the phosphorylation states of molecules involved in 14-3-3 signaling including PPAR signaling, IL-15 production, reelin signaling, and SSP compared to TBI alone (Figure 5G). Compared to controls, rTBI + 25-106 also evoked upregulation of PI3K/AKT, reelin, HIPPO, SSP, AMPK, 3-PI Deg, 3-PI biosynthesis (Bs), and SIPC signaling (Figure 5 Supplement 1A). Additionally, compared to controls, rTBI + 25-106 caused downregulation of molecules involved in HIPPO, 14-3-3, GLUR, HD, G-protein coupled receptor, AMPK, Reelin, and SSP signaling (Figure 5 Supplement 1B). Altogether, these data demonstrate the Cdk5/p25 signaling in the regulation of various neuronal pathways of injury involved in neuronal plasticity and synapse formation. Interestingly, this Cdk5 site enriched phosphoproteomic library derived from the acute phase of injury also unveiled the regulation of molecules implicated in chronic neurodegenerative conditions such as ALS and HD, two neurodegenerative conditions mediated via Cdk5 (Nguyen et al., 2001, Alvarez-Periel et al., 2018).

As an indication of how the molecules with increased or decreased phosphorylation states alter the biological processes within cells, gene ontology analysis using the open-source platform, Enrichr was employed. The 10 most significant biological processes altered by increased phosphorylation states following rTBI include those highly specified in structural and axonal functions, such as the regulation of microtubule polymerization/depolymerization, axonogenesis, cell morphogenesis in neuronal differentiation, neuronal projection morphogenesis, regulation of supramolecular fiber organization, and cellular component assembly (Figure 5H). This pattern of effects on structural and axonal processes was also observed from gene ontology analysis of downregulated phospho-sites following rTBI including regulation of microtubule polymerization/depolymerization, regulation of supramolecular fiber organization, axon development, cellular component assembly, and cell morphogenesis in neuron differentiation (Figure 5H). Upregulated phospho-proteins in rats subjected to rTBI and rTBI+25-106 also displayed similar patterns in the alterations of biological processes within structural and axonal gene ontologies including neuron projection morphogenesis, neuron projection development, cell morphogenesis/neuron differentiation, as well as, regulation of microtubule polymerization/depolymerization, regulation of dendritic spine morphogenesis, and synapse organization (Figure 5I). Rats subjected to rTBI + 25-106 also showed decreased phosphorylation in proteins associated with regulation of microtubule polymerization/depolymerization, regulation of supramolecular fiber organization, axon development, axonogenesis, and cell morphogenesis in neuron differentiation relative to rTBI alone animals (Figure 5I). The same pattern of structural and axonal processes was further observed in ontological analysis of rTBI + 25-106 rat hippocampi as compared to controls (Figure 5 Supplement 1C). Altogether, this study utilizing ontological analysis to prioritize potential downstream effects demonstrates broad alterations of the Cdk5 phospho-landscape within the hippocampus following rTBI.

### Total proteomics following rTBI

TBI may impart long-lasting impairments through alterations in protein expression that lead to cell death or inflammatory pathways ultimately leading to neuronal demise. To ascertain the effects of rTBI on protein levels as a net measure of the balance between expression and degradation, we conducted global proteomics of rTBI within the same samples prior to enrichment (Figure 5 Supplement 2). We noted 234 proteins demonstrated increased expression (FC ≥1.3) and 250 proteins with decreased expression (FC ≤ −1.3) (Figure 5 Supplement 2A) in rTBI hippocampus relative to controls. Additionally, we found 270 proteins exhibited increased expression (FC ≥1.3) and a decrease in 190 proteins (FC ≤ −1.3) in rTBI+25-106 hippocampus relative to rTBI alone (Figure 5 Supplement 2B). Also 321 proteins were increased (FC ≥1.3) and 261 proteins deceased (FC ≤ −1.3) in rTBI+25-106 vs control conditions (Figure 5 Supplement 2C).

The altered expression observed in the proteomic analysis within rTBI lysates was independently validated for two specific proteins (Figure 5 Supplement 2D-E). Proteomic analysis revealed animals subjected to rTBI displayed a 1.6 FC of Legumain (LGMN) protease following injury (Figure 5 Supplement 2A). Immunoblot analysis of hippocampal lysates confirmed LGMN is significantly upregulated in expression following injury (Figure 5 Supplement 2D). Additionally, Epoxide hydroxylase 2 (EPHX2) displayed a FC increase of 3.9 in rTBI rats compared to control rats (Figure 5 Supplement 2A). Similarly, immunoblot analysis displayed significant increases of EPHX2 in hippocampal lysates following injury (Figure 5 Supplement 2E). Altogether, these analyses reveal protein expression changes that may mediate aspects of rTBI.

## DISCUSSION

Rotational TBI remains the most clinically relevant form of brain trauma (Kleiven, 2013). However, understanding of these injuries has been, limited by the lack of rotational acceleration-induced injury models in rodents that accurately recapitulate patient phenotypes. Specifically, most preclinical rotational head trauma has been limited to large animal models due the conserved gyrencephalic brain structures lacking in rodent models (Armstead and Vavilala, 2020). While these large animal models of brain injury offer some physiologically relevant advantages, they remain costly and underpowered for genetic manipulations and preclinical screening of therapeutics (Sorby-Adams et al., 2018). The fundamental challenge with rodent models of rTBI has been to scale rotational forces to small brain structures. Helmet telemetry data in humans have determined a typical sports-related head trauma induces rotational accelerations between 1-3 Krad/s^2^ (Rowson et al., 2009, Rowson et al., 2012). Scaling to the 2 g rat brain, these equate to rotational forces greater than 350 Krad/s^2^ (Viano et al., 2009, Gennarelli et al., 1982). Previously, rodent models including the Medical College of Wisconsin (MCW) and Closed-Head Impact Model of Engineered Rotational Acceleration (CHIMERA) have provided insights into modeling these injuries in small mammals. These models have demonstrated histopathological and behavioral abnormalities following injury and show differential outcomes in these injury models in comparison to blast models of TBI (Stemper et al., 2016, Namjoshi et al., 2014, Sauerbeck et al., 2018). However, these models have remained limited largely to initial characterizations and, in some instances, lack equivalent scalar angular forces. Here, we validate our rTBI model’s rotational kinetics and demonstrate the feasibility for recapitulating numerous pathophysiological outcomes of head injuries across post-injury delays.

Several animal studies indicate neuroinflammation as a key TBI consequence (Shi et al., 2019). Depletion of infiltrating cells and halted neuroimmune responses improve behavioral and histological outcomes following cortical impact or fluid percussion injury in rodents (Willis et al., 2020, Witcher et al., 2021). Similarly, we show our rTBI model induces diffuse neuroinflammation. This inflammation is observed acutely (48 h) and persists into longer postinjury delay periods (7 & 14 d). We employed *in vivo* PET/CT imaging of TSPO as a potentially translational non-invasive diagnostic for rTBI. Increased TSPO expression likely indicates production of reactive oxygen species within neuroimmune cell populations such as microglia and astrocytes. This marker has been used clinically in patients with neurodegenerative conditions and in former professional football players exposed to chronic head injuries. The use of this marker as a diagnostic for acute brain injury in patients remains understudied (Coughlin et al., 2015, Werry et al., 2019). Interestingly, we observed no alteration in global brain uptake of TSPO, suggesting a mild or moderate neuroinflammatory phenotype. However, significant alterations in TSPO uptake were present when examining specific brain regions. The significance of region-specific neuroinflammatory responses such as these and the overall diagnostic sensitivity of this noninvasive technique patients with rTBI remains an important translational question.

Brain injury results in a multitude of cognitive and behavioral impairments. Some of the most common symptoms reported by TBI survivors are persistent impairments in cognitive abilities including deficits in memory, attention, and executive functioning (Arciniegas et al., 2002, Mathias and Mansfield, 2005). These features are commonly linked to structural damage in the hippocampal formation following injury (Tomaiuolo et al., 2004, Bigler et al., 2002), and numerous animal models point to impaired hippocampal-mediated contextual and spatial memory impairments following TBI (Sauerbeck et al., 2018, Arulsamy et al., 2018, Hernandez et al., 2018). In agreement, our rTBI model caused impaired hippocampal-dependent contextual fear memory and correlated with neuroinflammation in this region. Interestingly, amygdala-dependent cued fear memory was also impaired and exhibited a neuroinflammatory response. Structural damage to these areas, as well as brain-wide diffuse axonal injuries including interconnected efferent and afferent networks of the hippocampus/amygdala may be causally linked to these effects.

Underlying the observed memory impairment, rTBI caused an almost complete ablation of hippocampal long-term synaptic plasticity. This is consistent with the electrophysiological effects of other TBI models including fluid percussion, controlled cortical impact, and blast which also cause circuitry damage within the interconnected layers of the hippocampus (Goldstein et al., 2012, Perez et al., 2016, Titus et al., 2016). These alterations in hippocampal LTP occur in acute phases and persist into more chronic stages of TBI (Yousuf et al., 2016, Hernandez et al., 2018, Titus et al., 2016, Atkins, 2011). However, the long-lasting effects of rTBI on LTP remain to be further characterized.

TBI pathophysiology involves cellular depolarization, voltage-gated Ca^2+^ channel activation, massive glutamate release, and activation of Ca^2+^ -dependent proteases such as calpain (Werner and Engelhard, 2007, Lee et al., 2000). Calpain represents a potential therapeutic target for TBI (Saatman et al., 2010) and preclinical studies show promising calpain inhibitor neuroprotective activity (Buki et al., 2003, Czeiter et al., 2009, Kawamura et al., 2005, Meyer et al., 2014a). Cdk5/p25 activation is a principle downstream effector of excitotoxic calpain activation. Invocation of aberrant Cdk5 is broadly implicated in neuronal injury and neurodegeneration (Buki et al., 2003, Czeiter et al., 2009, Kawamura et al., 2005, Meyer et al., 2014a, Slevin and Krupinski, 2008, Pozo and J.A., 2016, Bk et al., 2019, Tsai et al., 2004, Wei and Tomizawa, 2007). Conditional knockout of Cdk5 is neuroprotective in models of ischemia and experimental brain injury (Yousuf et al., 2016, Meyer et al., 2014b). Transgenic overexpression of the aberrant co-activator of Cdk5, p25, induces rapid onset of Alzheimer’s disease in mice models (Cruz et al., 2003). Inhibition of p25 activity using small interfering peptides is neuroprotective in Parkinson’s disease models (Binukumar et al., 2015). Therefore, aberrant Cdk5 activity represents a critical therapeutic target for acute injury and neurodegenerative conditions.

There is currently no approved treatment to ameliorate the neuropathological consequences of TBI. Having recently discovered the first brain-active systemic Cdk5 inhibitor, 25-106 (Umfress et al., 2022), we tested it here as a possible treatment for rTBI. While use of 25-106 in the acute phase of rTBI appears promising, injury delay periods to emergency room treatments can vary between patients and nearly 80% of TBI patients are initially treated and then released from emergency departments (Capizzi et al., 2020). The efficiency of Cdk5 inhibition in more sub-acute and chronic stages of rTBI remains to be explored. Current TBI patient treatments are directed at symptom management including anticonvulsants for seizures, diuretics to reduce pressure in the brain, and stimulants and anti-anxiety medications to increase alertness and reduce negative mood states (Kirmani et al., 2016, Wakai et al., 2013, Levin et al., 2019, Silverberg and Panenka, 2019). Clinical trials have investigated numerous possible neuroprotective interventions including NMDA antagonists to reduce excitotoxicity and anti-inflammatory agents but have not significantly improved outcomes (Beauchamp et al., 2008, Bergold, 2016). On the other hand, clotting agents such as tranexamic acid can prevent brain edema in the acute phase, although this treatment is limited to intracranial hemorrhages and may not be a useful approach for repetitive mild injuries mediated by axonal injury and inflammation (Cap, 2019).

Given the efficacy of Cdk5 inhibition to block the deleterious behavioral and neuroinflammatory effects of rTBI, we used phosphoproteomics to enrich for phosphopeptides derived from potential Cdk5 substrates in the hippocampus. We report hundreds of novel potential Cdk5/p25 substrates phosphorylated in response rTBI but blocked when injury was followed by 25-106 treatment. Bioinformatic analysis implicated many of these in various cascades associated with cellular demise including AMPK, Ca^2+^, and neurodegenerative signaling. Importantly, human TBI survivors have a 56% increased risk of PD and a 2.3 times increased risk of developing AD (Gardner et al., 2018, Plassman et al., 2000). Since Cdk5/p25 activity is associated with the progression of both of these neurodegenerative conditions, it is likely that some of these substrates identified in the acute phases of rTBI likely contribute to longer lasting neuropathological effects of neurodegeneration. Understanding the dynamics of these phospho-substrates in the progression of TBI and as potential biomarkers of future brain degenerative conditions remains an exciting and expansive question generated by this work.

Several specific results within the global proteomic dataset derived here are also consistent with both preclinical models and clinical studies. For example, total proteome analysis in both rTBI and rTBI +25-106 showed the highest upregulation of bradykinin family members Kng1 (FC = 4.0, rTBI FC = 6.6 rTBI + 25-106) and Kng2 (FC = 3.8, rTBI FC = 6.0 rTBI + 25-106). These proteins have been heavily implicated in animal TBI models as well as clinical studies (Trabold et al., 2010). Kng1/Kng2 serve as ligands for B2 bradykinin receptors. In alignment with these targets as mediators of rTBI, B2 receptor antagonists have been tested in clinical trials for TBI treatment, (Beauchamp et al., 2008). We note that the induction of this inflammatory signaling cascade was not responsive to 25-106, and thus may exemplify mechanisms of injury that are not necessarily effectors of aberrant Cdk5 activation. Altogether, this work brings forth a new model to the field of brain injury, assesses a novel neuroprotective treatment, and progresses our knowledge of the mechanisms mediating neuronal death caused by head trauma and other excitotoxic conditions.

## MATERIALS AND METHODS

### Animals

Male Sprague Dawley rats (9-10 months old, Envigo) were used for all TBI experiments. All were single housed and maintained on a normal 12 h day/night cycle. All experiments were performed under approved protocols by the University of Alabama at Birmingham (UAB) Institutional Animal Care and Use Committees (IACUC).

### Rotational Traumatic Brain Injury (TBI) procedure

All rats were habituated to the room in which brain injuries occurred for 1 h, anesthetized using 1.5% isoflurane vapor, and sedation was maintained throughout the procedure via a nose-cone within the subject’s helmet. Anesthetized rats were placed in a custom designed harness that was inserted into the base of the pendulum and locked tightly into place. The rat head was fitted into a freely rotational helmet with foam padding to secure head placement and rotation in one direction. Flowing N_2_ gas was allowed to fill an Arduino controlled closed solenoid value connected to an air motor and the pendulum was adjusted to a 90° angle raised approximately 84 cm from the base of the machine. Launch was electronically executed using custom software (LabVIEW) to open the solenoid allowing N_2_ to propel the pendulum forward towards the ventral strike plate. Helmet impact results in a 90° rotation of the helmet and animal’s head in both directions. Rotational forces were derived in LabVIEW from helmet velocity and exported into Excel. Each animal received 10 repetitive impact rotations over a 30 min period. Following injury each animal was assessed for normal locomotion and righting reflex abilities.

### PET/CT imaging

Noninvasive positron emission tomography-computed tomography (PET/CT) imaging was conducted 7d post TBI with [^18^F]DPA-714, a translocator protein (TSPO) radioligand produced at the Cyclotron Facility at the University of Alabama at Birmingham. Rats were injected with 300 μCi (11.1 MBq) of [^18^F]DPA-714 intravenously. Rats were immediately scanned using a GNEXT small animal PET/CT (Sofie Biosciences) for 30 min. Imaged regions of interests (ROIs) within the brain were drawn with CT guidance using VivoQuant (Invicro) and a 3D rat brain atlas (Allen brain atlas) was applied to automatically segment each brain region. The mean, maximum, hotspot, and heterogeneity of standard uptake values (SUVs) were determined using the formula: SUV = [(MBq/mL) x (animal wt. (g))/injected dose (MBq)].

### Histology

Histology was performed as described (Umfress et al., 2021). Briefly, rats were transcardially perfused with 1X PBS/50 mM NaF followed by 10% formalin fixation. Brains were submerged in 10% formalin and fixed overnight, paraffin embedded and serially sectioned at 5 μm. Sections were deparaffinized, rehydrated, and subjected to citrate antigen retrieval. Sections were permeabilized and blocked in 0.03% Trition X-100/PBS containing 3% goat serum. Immunohistochemistry was performed using glial fibrillary acidic protein (GFAP) (1:1000; Millipore) and ionized Ca^2+^-binding adapter protein (1:1000; Wako) incubated in 0.3% Tween 20/PBS. Fluorescent visualization was performed using secondary antibodies Cy3 anti-rat (1:500) and Alexa 488 anti-mouse (1:200) (Jackson Immuno) and imaged using an Olympus BX60 microscope mounted with Olympus DP74 digital camera. All stains were performed using a minimum of 3 sections per animal and 4-6 animals per group. All stains within groups were analyzed using the same background thresholds (0-72; GFAP) and (4-110; IBA1). Percent area of positive stain was determined using ImageJ software (Fiji), and stains within the same animal were averaged to obtain final values.

### *Ex vivo* acute brain slice pharmacology

Brain slice pharmacology was performed as described (Umfress et al., 2022). Briefly, brains were rapidly decapitated and submerged in ice cold Normal Kreb’s solution (125 mM NaCl, 2.5 mM KCl, 1.25 mM NaH_2_PO_4_, 25 mM NaHCO_3_, 1.1 mM MgCl_2_, 2 mM CaCl_2_ and 25 mM glucose). Brains were coronally sectioned at 350 μm using a vibratome in regions of interest. Slices were recovered in oxygenated Krebs solution at 30°C. Following recovery, slices where incubated in Kreb’s containing pharmacological interventions indicated of NMDA/Glycine (100 μM NMDA, 50 μM Gly, 1h), 25-106 (10 μM, 1 h). Following treatment, slices were snap frozen in dry ice to terminate treatment. For Oxygen-Glucose Deprivation (OGD) slice studies, following recovery, slices were incubated in a deoxygenated (N_2_ bubbled in solution 1 h) Kreb’s solution substituting sucrose for D-glucose (125 mM NaCl, 2.5 mM KCl, 1.25 mM NaH_2_PO_4_, 25 mM NaHCO_3_, 1.1 mM MgCl_2_, 2 mM CaCl_2_ and 25 mM sucrose) for 30 min. Following OGD treatment, slices were incubated in 0.125% 2,3,5-Triphenyltetrazolium chloride (TTC) viability stain for 20 min. Stained slices were fixed in 4% PFA for 10 min and scanned. TTC viability was assessed by mean intensity of staining and expressed as percentage of control viability.

### Primary Neuronal Cell Culture and Oxygen-Glucose Deprivation

Primary cortical neuronal cultures were derived from mouse embryos on embryonic day 15 (E15) as previously described (Hossain et al., 2021). Briefly, pregnant dams were sacrificed, embryonic cortices were dissected and dissociated in medium consisting of DMEM+ 20% horse serum. Isolates were washed in serum free medium and trypsin digested at 37°C. Cortices were subsequently triturated in neurobasal media + 10 mM glucose, 1 mM GlutaMax,1 mM sodium pyruvate, and B-27, strained and plated at a density of 5 × 10^5^ cells/ml on cell culture plates. On the day in vitro (DIV) 2, the cultures were treated with 40 μM 5-fluoro-2-deoxyuridine Mature neurons DIV 11–12 for oxygen glucose deprivation (OGD) studies. To induce OGD, primary neurons where incubated in OGD buffer (NaCl 116 mM, KCl 5.4 mM, MgSO_4_ 0.8 mM, NaHCO_3_ 26.2 mM, NaH_2_PO_4_ 1 mM, CaCl_2_ 1.8 mM, glycine 0.01 mM, pH 7.4) pre-bubbled with OGD gas (5% CO_2_, 10% H2 and 85% N_2_) in a hypoxia chamber at 37°C, connected to an O_2_ sensor/monitor (Biospherix Ltd. NY, USA) for 90 min. The O_2_ was maintained <1% for the duration of treatment. OGD was terminated by reperfusion of normal neuronal media and restoration of normoxic conditions. Pharmacological interventions included treatments with 10 μM 25-106 or 0.01 % DMSO control at the onset of OGD induction. Cell viability was assessed 24 h after OGD via Alamar blue assay. Viability was expressed as a percentage of live cells as compared to control (no OGD) neurons.

### Immunoblotting

Immunoblotting was performed as previously described (Umfress et al., 2021). Briefly, rat brains were rapidly dissected and submerged in a 4°C solution of PBS/50 mM NaF. Subregions were dissected and snap frozen. Brain regions were subsequently homogenized in 1% SDS/50 mM NaF and sonicated at 40 dB pulses until tissue was completely homogenized. Protein concentrations were determined from lysates via BCA protein assay. Samples were diluted in 4x lysis buffer and proteins were separated by molecular weight via SDS-PAGE. Proteins were then transferred to 0.45 nm nitrocellulose, blocked in Licor Blocking Buffer, and incubated with 1°antibody (Ab) overnight. Membranes were washed with 1xTBS-T and incubated with Licor fluorescent 2° Ab for 1 h at room temperature. Proteins expression was visualized using Licor Odyssey CLx membrane scanner. Arbitrary units of florescent intensity for each protein band was quantified using ImageStudio. Phospho-band intensity was normalized to total protein bands, total protein bands were normalized to actin loading controls, and cleavage products were normalized to un-cleaved total protein. Antibodies used include p35/p25 (Cell Signaling Technology), phospho-Ser549 and total Synapsin I (PhosphoSolutions), Fodrin (Enzo Life Sciences), phospho-Thr75- and total DARPP32 (Cell Signaling Technology), EPHx2 (Abcam), Legumain (Cell Signaling), Actin (Invitrogen).

### Neurophysiological Recordings

Neurophysiological studies were conducted in rats 7 d post injury as described (Hernandez et al., 2018). Briefly, brains were rapidly dissected in ice-cold artificial cerebrospinal fluid (ACSF; 75 mM sucrose, 87 mM NaCl, 2.5 mM KCl, 1.25 mM NaH_2_PO_4_, 25 mM NaHCO_3_, 7 mM MgCl_2_, 0.5 mM CaCl_2_ and 10 mM glucose). Transverse hippocampal slices (350 μ) were sectioned using a vibratome (Leica Microsystems Inc., VT1000S) in NMDG cutting/recovery solution (N-methyl D-glucamine (100 mM), KCl (2.5 mM), NaH_2_PO_4_ (1.2 mM), NaHCO_3_ (30mM), HEPES (20 mM), MgS0_4_ (1 0mM), CaCl_2_ (0.5 mM), and glucose (25 mM) at 30°C (pH 7.3-7.4). After 2 min, slices were transferred to HEPES holding solution NaCl (92 mM), KCl (2.5 mM), NaH_2_CO_3_ (30 mM), NaH_2_PO_4_ (1 mM), HEPES (20 mM), D-Glucose (25 mM), MgCl_2_ (1 mM), CaCl_2_ (1 mM) for 1 h at 30°C. Slices were allowed to incubate for 30 min in recording solution of oxygenated Kreb’s (125 mM NaCl, 2.5 mM KCl, 1.25 mM NaH_2_PO_4_, 25 mM NaHCO_3_, 1.1 mM MgCl_2_, 2 mM CaCl_2_ and 25 mM glucose) prior to recording.

Recordings were performed using a Multiclamp 700A amplifier with a Digidata 1322 and pClamp 10 software (Axon, Molecular devices, LLC). Field excitatory postsynaptic potentials (fEPSP) from CA1 were evoked by square current pulses (0.1 ms) at 0.033 Hz with a bipolar stimulation electrode (FHC, Bowdoinham, ME). Stimulus intensity was defined using a stimulus intensity required to induce 50% of the maximum EPSP slope using the input-output curves. The sample intensity was used for PPR recordings across different intervals. A stable baseline was recorded for at least 10 min prior to high frequency stimulation (HFS, 4 trains, 100 Hz, 1 s duration, separated by 20 s). Post-tetanic potentiation (PTP) was analyzed by taking the average of the slopes from the traces recorded during the first 2 min after HFS. LTP was assessed for at least 45-50 min following HFS. The PPR values were calculated by dividing the second fEPSP slope by the first fEPSP slope (fEPSP2/fEPSP1). All recordings were performed in the absence of any drug treatment and only 1 or 2 slices were recorded from each individual rat. Data were analyzed with Clampfit 10 software (Axon, Molecular devices, LLC).

### Neurobehavior

Fear conditioning studies were performed as described (Plattner et al., 2015). Briefly, rats were habituated to the behavioral room for 1 h before experimentation. Rats were placed in fear conditioning chambers (Med Associates) to establish baseline freezing rates between cohorts. The following day rats were subjected to fear conditioning training. Each rat was allowed to freely explore the chamber for 2 min followed by a 30 s tone terminating in a mild foot shock (0.7 mA). Rats remained in the chamber 2 min after shocking before returning to their home cage.

Context-dependent fear memory was assessed 24 h post-shock training. Rats were reintroduced into the conditioning box for 5 min. Freezing responses (motionless except respirations) were recorded using VideoFreeze software. Cued fear conditioning memory was assessed 27 h post-shock training where rats were allowed to explore a novel context with novel odor (vanilla). The rats were left exploring the novel context for 3 min without tone followed by a 3 min period with the training tone playing. Freezing responses (motionless except respirations) were recorded.

Shock sensitivity and nociceptive responses were assessed by returning rats to fear conditioning chambers and evaluating minimal thresholds to induce rat flinching, jumping, and vocalization of pain across adverse stimuli (0-1.5 mA) shocks.

### Discovery proteomics

Freshly dissected and snap frozen rat hippocampi were sonicated, centrifuged, reduced with DTT, and alkylated with iodoacetamide. Total protein for each sample (10 mg) was trypsin digested, purified over C18 columns (Waters), enriched using the PTMScan Phospho CDK + CDK/MAPK Substrate Motif Antibodies (#9477/#2325 Cell Signaling Technology) and purified over C18 tips as previously described (Stokes et al., 2015). For total proteome analysis an additional 100 mg of each sample was digested with LysC and trypsin and digested samples were purified over C18 tips, labeled with TMT 10-plex reagent (Thermo), bRP fractionated (96 fractions concatenated non-sequentially to 12), and C18 purified for LC-MS/MS as previously described (Possemato et al., 2017). LC-MS/MS analysis was performed using an Orbitrap-Fusion Lumos Tribrid mass spectrometer as described (Possemato et al., 2017, Stokes et al., 2015) with replicate injections of each sample run non-sequentially for the phosphopeptide analysis. Briefly, peptides were separated using a 50 cm x 100 μM PicoFrit capillary column packed with C18 reversed-phase resin and eluted with a 90 min (Phospho) or 150 min (TMT total proteome) linear gradient of acetonitrile in 0.125% formic acid delivered at 280 nl/min. Full MS parameter settings are available upon request. MS spectra were evaluated by Cell Signaling Technology using Comet and the GFY-Core platform (Harvard University) (Eng et al., 2013, Huttlin et al., 2010, Villen et al., 2007). Searches were performed against the most recent update of the NCBI Rattus norvegicus database with a mass accuracy of +/-20 ppm for precursor ions and 0.02 Da product ions.

Results were filtered to a 1% peptide-level FDR with mass accuracy +/-5 ppm on precursor ions and presence of a phosphorylated residue for CDK Substrate enriched samples. TMT total proteome results were further filtered to a 1% protein level false discovery rate. Site localization confidence was determined using AScore (Beausoleil et al., 2006). All CDK Substrate quantitative results were generated using Skyline (MacLean et al., 2010) to extract the integrated peak area of the corresponding peptide assignments or in GFY-Core using signal: noise values for each peptide and summing individual signal: noise values for all peptides for a given protein for TMT total proteome data. Accuracy of quantitative data was ensured by manual review in Skyline or in the ion chromatogram files. Quantitative data was normalized across samples using median abundance for CDK Substrate data or sum signal: noise for TMT total proteome data.

### Pathway and Ontology Analysis

Differentially modified phosphoproteins across all groups upregulated ≥1.3-fold change or downregulated ≤ −1.3 -fold change were subjected to Ingenuity pathway analysis (IPA), (Qiagen). Log-transformed p-values of significantly enriched canonical pathways and the phosphorylated/dephosphorylated molecules comprising each pathway were used to construct dot plots of canonical pathways significantly Up/downregulated in each condition. IPA Z-scores were used to predict activation or inhibition of each canonical pathway. Phosphoproteins exhibiting fold changes of greater ≥ 1.3 or ≤ −1.3 were subjected to gene ontology enrichment analysis via open source gene enrichment analysis software, Enrichr (Chen et al., 2013). Biological processes and molecular functions from each gene list were derived based and the top 10 processes/functions were derived based on adjusted p-values.

### Statistical Analysis

Prior to analysis, all data was examined for normalcy using the Shapiro-Wilk test. In cases of parametric data with normal distributions of two group means Student’s *t*-test were used. In cases of non-parametric comparison of means, the Mann-Whitney test was employed. When comparing more than two group means, one or two-way ANOVAs were used with Holm-Sidak post-hoc tests. When comparing two experimental variables within the same animal, Two-way repeated measures ANOVA was used. For all data, * = p < 0.05, ** = p < 0.01, ** = p < 0.001, *** = p < 0.0001. All statistical analysis was performed using Prism 6 (GraphPad Software, Inc.)

## ACKNOWLEDGEMENTS

We thank the UAB Behavioral Assessment Core for assistance. We thank UT Dallas Mechanical Engineering Senior Design Program and the students, Carlo Wayan, Curtis Anderson, Jose Becerra, Stacy Donovan, and Athanasios Spiropoulos for the design and construction of the model. We thank Divya Shaw and the UT Arlington Mechanical and Aerospace Engineering Dept., and Naveen Varma and the UAB School of Engineering for help with model development. We thank V. Yang for assistance with data library deposition. The authors acknowledge the UAB Comprehensive Cancer Center’s Preclincal Imaging Shared Facilitity (P30CA013148). We thank the Dept of Psychiatry and Carol Tamminga for support of this work. We appreciate the IACUC staffs at UTSW and UAB for help in developing this model.

## AUTHOR CONTRIBUTIONS

A.U., A.C., S.P.S.D., D.E., R.A., A.M., A.S., I.H., S.A., T.W., C.S., and M.S. conducted experiments, collected data, and performed formal analysis. H. Luo. and H.Lu. engineered the model. S.S. and A.N. synthesized reagents. D.C., N.K., and S.M. conducted analysis and data visualization. A.U., C.R., J.A.B. conceived the study and experimental design.

All authors contributed to editing and approval of the final manuscript

