## Supplementary Material for "Characterization and Preclinical Treatment of Rotational Force-Induced Brain Injury"

Umfress, et al. Figure 1 Supplement 1

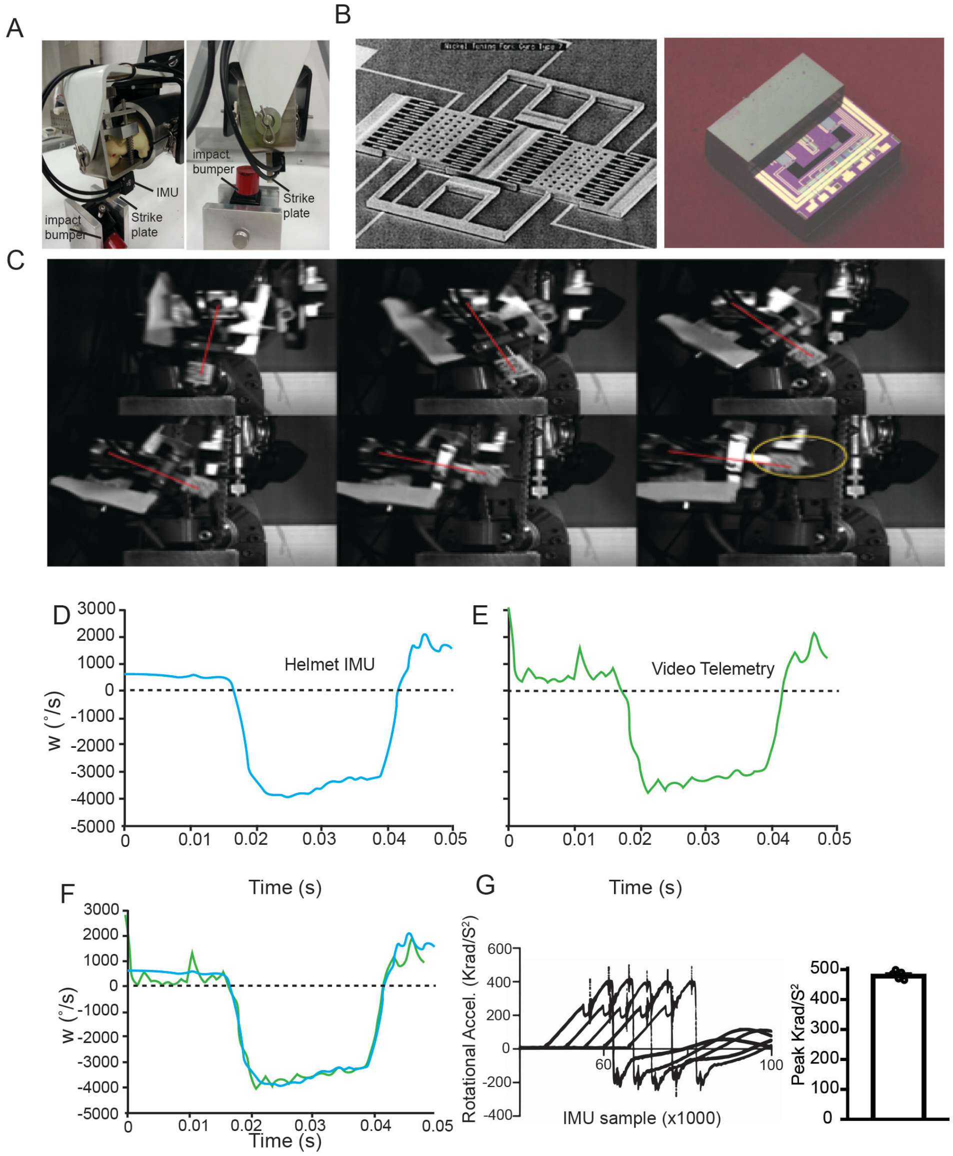


**Figure 1 Supplement 1. rTBI Model validation and kinetics.** A. Side view of helmet assembly and restraint with artificial rat. Note Model 633 IMU mounted on helmet strike plate below rat head with front view showing strike plate and impact bumper B. Model 633 6DOF inertia measurement unit. Oscillating variable capacitance gyro sensor combs (right) and half cover removed view of IMU showing acceleration sensor, mass, supports, and circuitry. C. Temporally sequential high- speed camera recording frames used to derive acceleration rates. D. Sample recording from IMU. E. Video telemetry calculated rate vs. time. F. Overlay of helmet IMU and Video telemetry angular speeds ($^{o}$/S) as a function of time. G. Labview derived rotational accelerations (Krad/S$^2$) of 5 trial runs (left). Peak rotational accelerations for each run with standard error (right).

Umfress, et al. Figure 2 Supplement 1

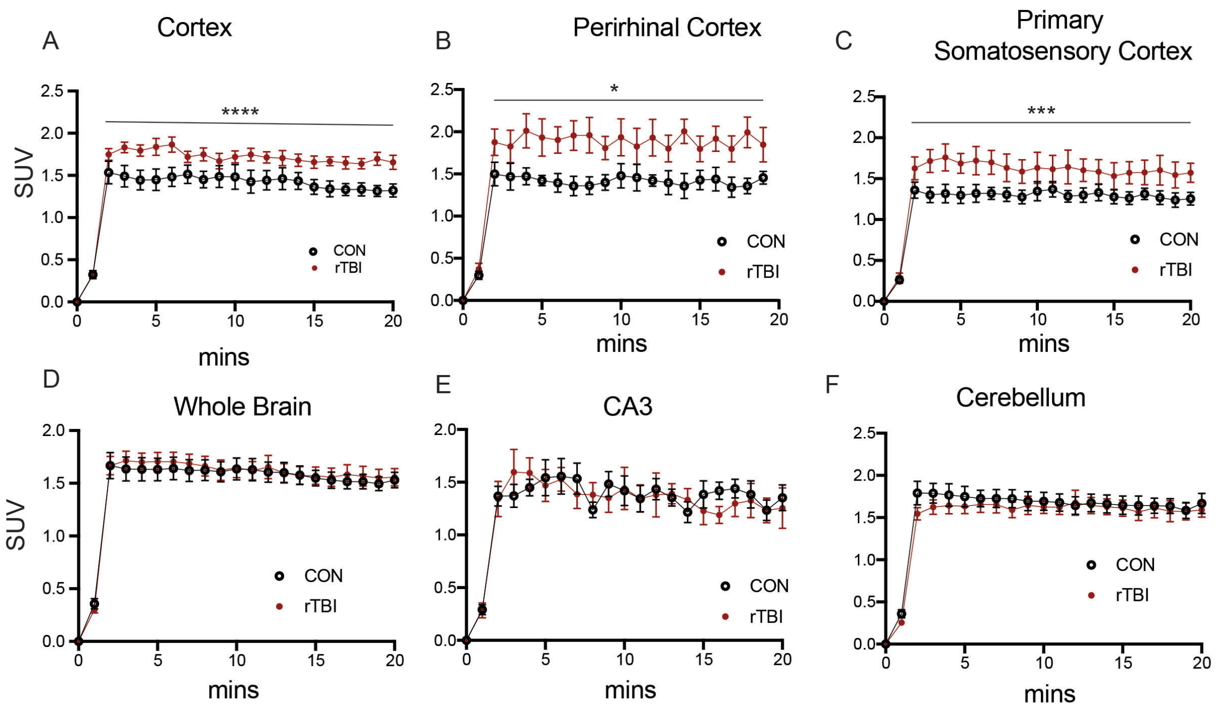


**Figure 2 Supplement 1. PET/CT analysis of diffuse TSPO uptake into the brain** Standard uptake values in A. cortex (Time: F(20,200 ) = 197.0, p < 0.0001; Treatment: F (1, 10) = 5.580, p = 0.0398; Interaction: F(20,200) = 3.063, P < 0.0001) two-way-RM ANOVA B. Perirhinal cortex (Time: F(20,200) = 61.32, p= 0.0130; Treatment: F(1,10) = 6.020, p = 0.0341; Interaction: F(20,200) = 1.917, p = 0.0130) C. Primary somatosensory cortex (Time: F(20, 200) = 132.4, p < 0.0001; Treatment: F(1,10) = 3.418, p = 0.0942; Interaction: F(20,200) = 2.374, p = 0.0013). D. whole brain (Time: F(20,200) = 402.6, p<0.0001; Treatment: F(1,10) = 0.06724, p = 0.8007; Interaction F(20,200) = 0.8649 E. CA3 (Time: F(20,200) = 47.91 p < 0.0001; Treatment: F(1,10) = 0.02460, p=0.8785; Interaction F(20,200) =1.030, p = 0.4283). F. Cerebellum (Time: F(20,200) = 277.4 p < 0.0001; Treatment F(1,10) = 0.3043 p = 0.5933; Interaction F(20,200) = 1.255, p = 0.2138. All data are means ± SEM, *p < 0.05, *< 0.01, ***< 0.001. All statistical analysis consisted of two way-RM ANOVA.

Umfress et al Figure 4 Supplement 1

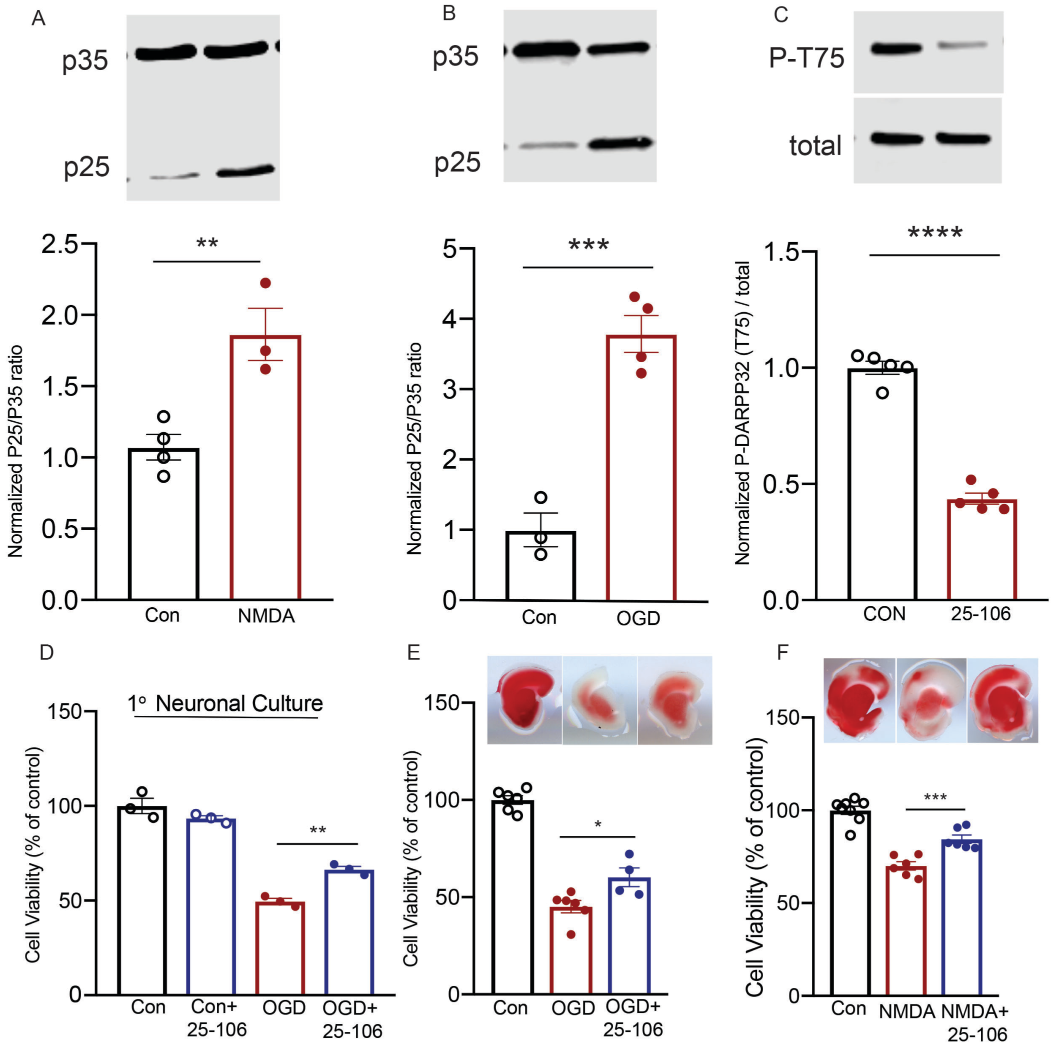


**Figure 4 Supplement 1. Aberrant Cdk5/P25 activity mediates excitotoxic cell death.** A. quantitative immunoblot of acute brain slices for p35/25 after NMDA/Glycine (100/50 μM, NMDA/Gly) treatment, $p = 0.0081$ Student's *t*-test B. Quantitative immunoblot of acute brain slices for p35/25 after OGD treatment, $p=0.0007$ Student's t-test. C. Quantitative immunoblot of acute brain slices for phospho-Thr75 DARPP32 after 1 h treatment 25-106 (10 μM), $p < 0.0001$ Student's *t*-test. D. Alamar Blue viability assay of primary neuronal culture following OGD treatment and preincubation with 25-106 $F(3,8)= 95.38$, $p < 0.0001$ ANOVA, OGD-OGD+ 25-106 $p = 0.0022$, Holm-Sidak post hoc. E. TTC viability staining of acute brain slices subjected to OGD following preincubation with 25-106 $F(3,13) = 84.60$ $p < 0.0001$ ANOVA (OGD vs OGD +25106, $p = 0.0080$, Holm-Sidak *post hoc*). F. TTC viability staining of acute brain slices subjected to NMDA/Gly (100/50 μM) treatment following preincubation with 25-106 $F(2,17) = 45.19$ $p < 0.0001$ ANOVA (NMDA vs NMDA + 25-106, $p= 0.0005$, Holm-Sidak *post hoc*). All data are means ± SEM, *p < 0.05, **P<0.01, ***P<0.001 ****P<0.0001.

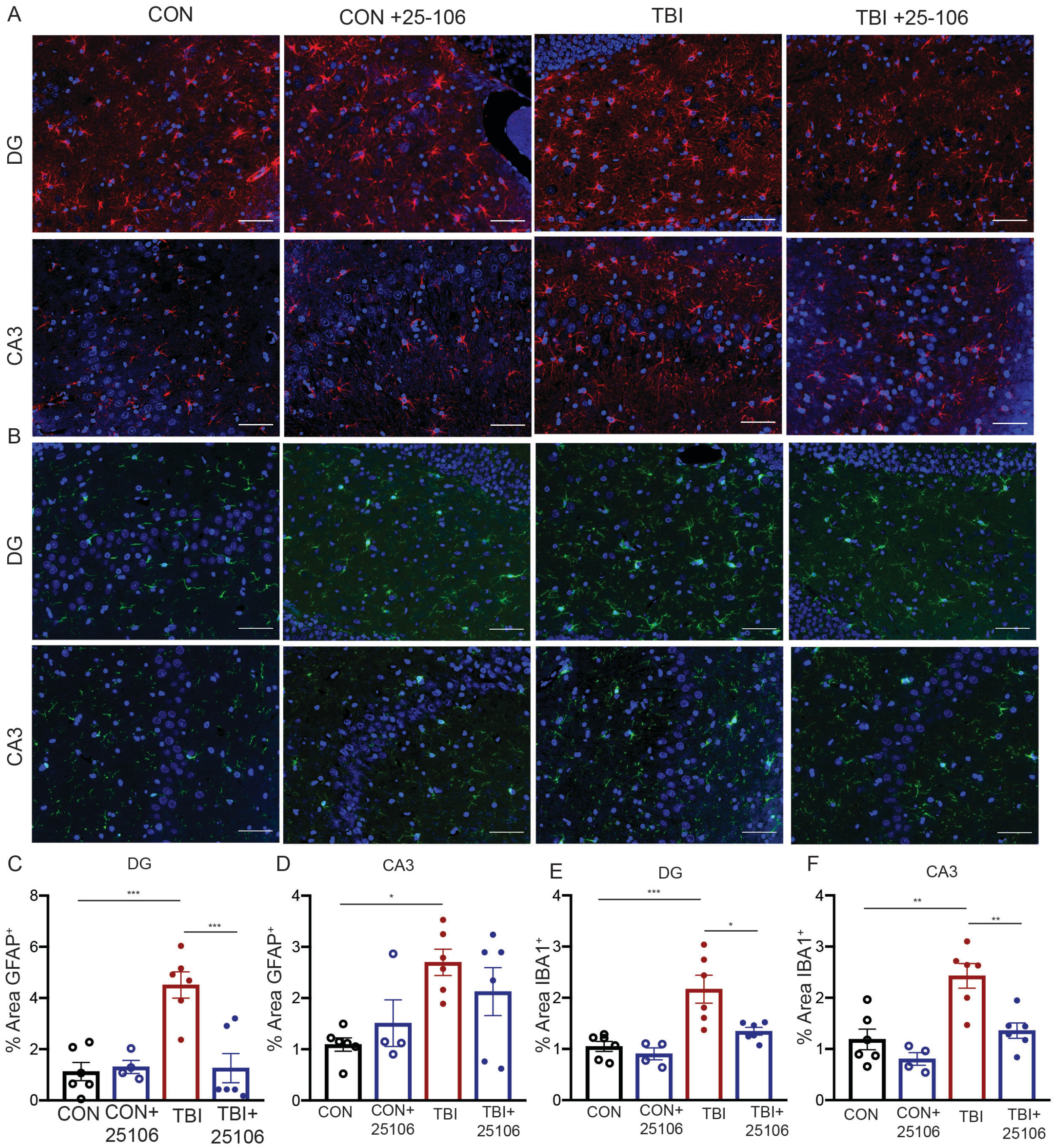


**Figure 4 Supplement 2. Cdk5 inhibition provides *in vivo* neuroprotection.** Immunohistochemical staining and quantitation of GFAP+ astrocytes within the DG (A, C) $F(3,18) = 12.74$, $p = 0.0001$ ANOVA and CA3 (A, D) $F(3,18) = 4.494$, $p = 0.0160$ ANOVA subfields of the hippocampus. Immunohistochemical staining and quantitation of Iba1+ microglia within the DG (B, E), $F(3,18) = 11.11$, $p = 0.0002$ ANOVA and CA3 (B, F), $F(3,18) = 12.33$, $p = 0.0001$ ANOVA subfields of the hippocampus. All data are means ± SEM, $p < 0.05$, $^{**}<0.01$. Scale bars = 100 μm.

Umfress, et al. Figure 5 Supplement 1

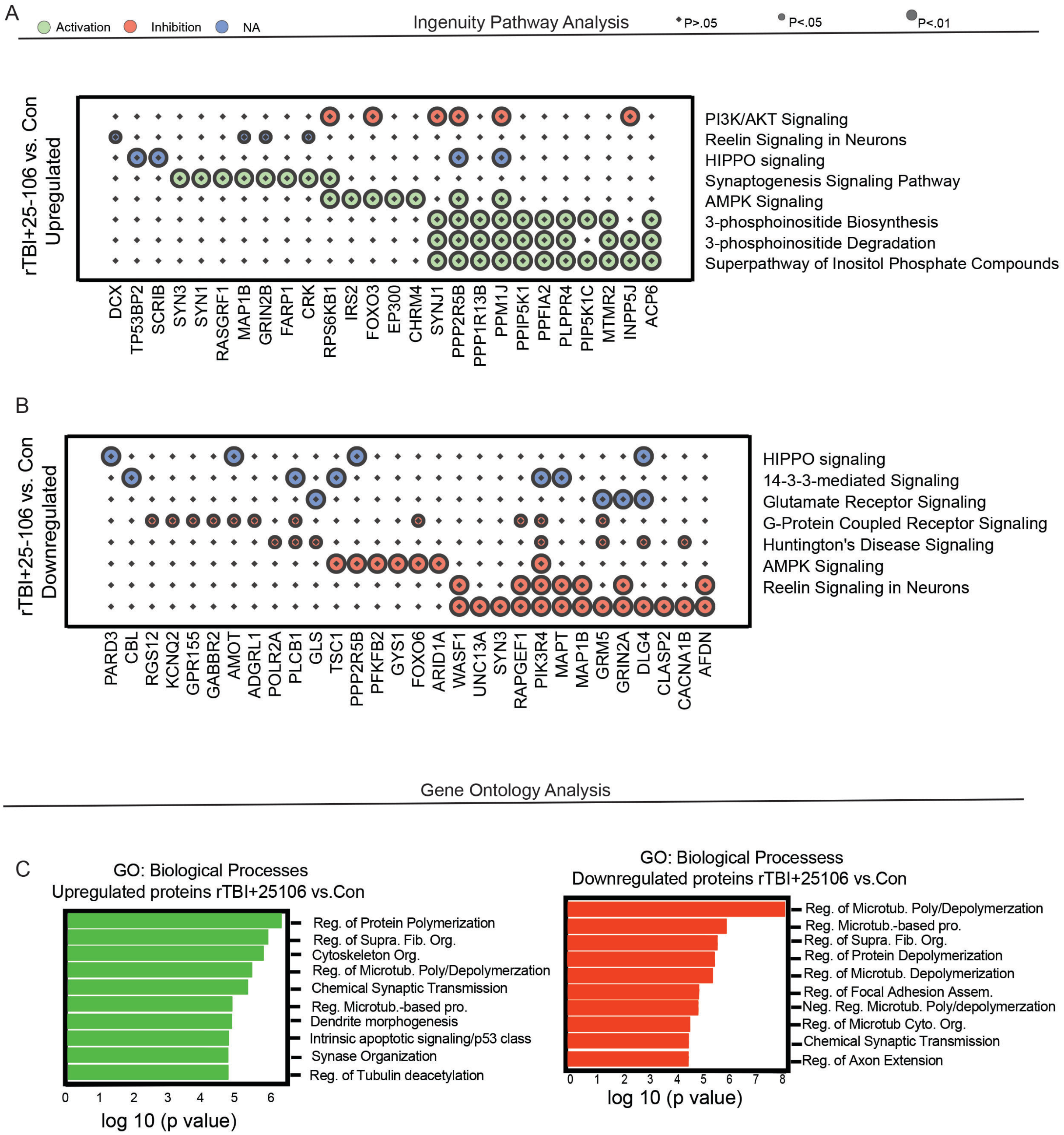


**Figure 5 Supplement 1. Pathway and ontological analysis of rTBI+25-106 vs Con rats.** A. Dot-plot display of Ingenuity Pathway Analysis of phospho-proteins and canonical pathways upregulated FC≥1.3 by rTBI + 25106:Con (Top) and B. downregulated with a FC ≤ -1.3 by rTBI + 25-106:Con (bottom). C. Gene ontology of phosphoproteins biological processes upregulated FC≥ 1.3 (left, green) or downregulated FC ≤ -1.3 (right, red) in rTBI + 25-106:Con.

Total proteomics of Injury

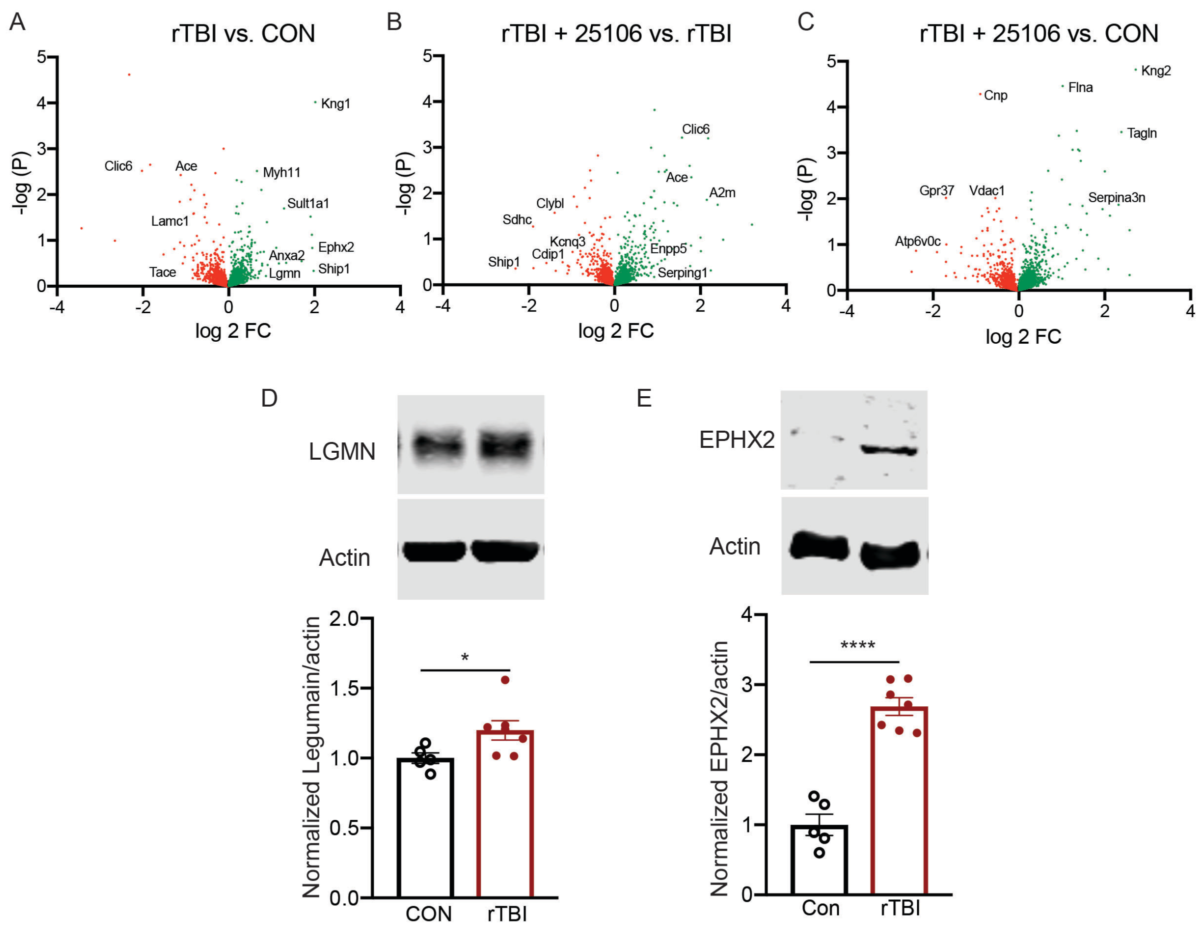


**Figure 5 Supplement 2. Total proteomics and validation.** A. Volcano plot of differentially expressed total proteins between rTBI and con rats (increased sites in green decreased sites in red). B. Volcano plot of differentially expressed proteins between rTBI + 25-106 and rTBI alone rats. C. Volcano plot of differentially expressed proteins between rTBI + 25-106 and con rats. D. Quantitative immunoblot for protein expression ratio of LGMN/ actin, $p = 0.0494$ Student's t-test. E. Quantitative immunoblot for protein expression ratio of EPHX2/ actin, $p < 0.0001$. Data are means ± SEM, *$p < 0.05$, ****< 0.0001.
